# Hotter chili peppers by hybridisation: heterosis and reciprocal effects

**DOI:** 10.1101/2021.09.16.460669

**Authors:** Emmanuel Rezende Naves, Federico Scossa, Wagner L. Araújo, Adriano Nunes-Nesi, Alisdair R. Fernie, Agustin Zsögön

## Abstract

The selection of pure lines followed by crosses to create superior hybrids is one of the oldest strategies in crop breeding. However, in domesticated species of the *Capsicum* genus hybrid breeding has lagged, in part due to a lack of detailed information about the phenotypic and metabolic consequences of hybridization. Here, we performed reciprocal crosses between four inbred varieties of two species of cultivated *C. chinense* (cv. Habanero and Biquinho) and *C. annuum* var. *annuum* (cv. Jalapeño and cv. Cascadura Ikeda). These varieties were specifically selected for their highly divergent traits, including plant growth habit, fruit size, shape and pungency: Habanero and Jalapeño peppers are highly pungent forms, particularly popular in Mexico. The Biquinho cultivar of *C. chinense* and the Cascadura Ikeda bell pepper are traditional sweet cultivars from Brazil. From the parental genotypes and from the progeny of the reciprocal crosses, we measured 28 phenotypic traits, including plant growth, and yield, 32 fruit shape parameters, and 50 fruit pericarp and placenta metabolites, including capsaicinoids. We found large differences for agronomic and metabolic traits between the genotypes, including heterosis for pungency and reciprocal effects for agronomic traits. We also show that the strong association between fruit shape and pungency can be broken in intraspecific hybrids, paving the way for the precision breeding of novel varieties.

**Once sentence summary:** Hybrids of Capsicum peppers display heterosis and reciprocal effects for agronomic traits and fruit pungency

## Introduction

Heterosis, or hybrid vigour, is a genetic phenomenon whereby the hybrid progeny of different inbred strains, races or species of plants or animals results in increased vigour or performance. Charles Darwin described the phenomenon in the late 19^th^ century, but it was only in 1914 that G. H. Shull coined the term ‘heterosis’ and properly framed it conceptually (Hochholdinger and Baldauf, 2018). Large-scale cultivation of hybrid crops was introduced in the 1930s and has been growing steadily since, providing yield increases of up to 50%, depending on the crop. Although the genetic, epigenetic and expression bases for heterosis have been explored extensively (Chen, 2013), the metabolic aspects behind this phenomenon remains obscure. Significant advances have been made in understanding the metabolic basis of heterosis in maize (Riedelsheimer et al., 2012; Lima et al., 2017; Li et al., 2020) with the aim of establishing a ‘multi-omics’ platform for predictive hybrid breeding (Westhues et al., 2017). However, a similar effort is lacking in horticultural species, even though hybrid breeding is extensive in these crops (Yu et al., 2021). Here, we explore agronomic and metabolic aspects of heterosis in hybrids of peppers (*Capsicum* spp.).

Among the 41 species described in the *Capsicum* genus (Barboza et al., 2019), only five (*Capsicum annuum* var. *annuum, C. frutescens, C. chinense, C. baccatum* var. *pendulum*, and *C. pubescens*) have been domesticated (Pickersgill, 1997). With the exception of *C. pubescens*, which has limited importance today as a cultivated species, all remaining domesticates have been specifically bred to give rise to a high diversity of cultivars (Carrizo García et al., 2016), providing an excellent resource for the study of phenotypic variation induced by artificial selection (Pickersgill, 2018). The selection for fruit-related traits has brought in an amazing diversity of fruit colors, size and shapes (Scossa et al., 2019), including a wide variation of pungency levels, with cultivated varieties covering the full spectrum from completely sweet to prohibitively hot fruits (Muñoz-Ramírez et al., 2018). The pungent taste of peppers is conferred by a class of vanillylamides, called capsaicinoids, which accumulate during ripening in the placenta of hot varieties (Naves et al., 2019). Capsaicinoids and other pepper phytochemicals have potential uses in the agrifood, cosmetic and pharmaceutical industries as replacements for synthetic additives (Baenas et al., 2019).

Being an autogamous species (with the only exception of *C. cardenasii*, which is an obligate outcrosser (Pickersgill, 1997), and as a consequence of the continuous process of purging deleterious mutations through artificial selection, cultivated varieties of *Capsicum* do not show inbreeding depression. However, heterosis (or hybrid vigour) has not been investigated in detail in *Capsicum* breeding, but increased yield, biomass and pungency was shown in some selected hybrids of *C. annuum* (Garcés-Claver et al., 2007; Singh et al., 2014). It has recently been shown that alterations in gene expression profile are associated with heterosis in pepper hybrids (Yang et al., 2021). Cultivation of F_1_ hybrids rather than varieties could lead to higher yield and fruit quality, although many traits of agricultural importance are strongly influenced by environmental factors (Tripodi et al., 2020). Hybridization breeding also has the advantage of combining desirable horticultural and resistance traits faster than conventional pure line and pedigree selection, as it allows the combination of dominantly inherited traits (Zhao et al., 2015). However, the commercial exploitation of hybrids, which instead represent the fundamental breeding form in other crops, has not found so far large application in *Capsicum*, due to the existence of incompatibility barriers between species (Onus and Pickersgill, 2004) and to the absence of a convenient male-sterility system to avoid self-fertilization (Kim and Zhang, 2018).

*Capsicum* breeding programs have been limited by the relatively narrow genetic base exploited, mostly within the *C. annuum* cultivars (Pereira-Dias et al., 2019). Even though F_1_ hybrids are widely grown for commercially produced sweet pepper, this is not the case for hot peppers. Here, we performed an in-depth analysis of the developmental, agronomic, and metabolic consequences of hybridization between of pungent and non-pungent commercial cultivars of *C. chinense* (cv. Habanero and cv. Biquinho) and *C. annuum* var. *annuum* (cv. Jalapeño and cv. Cascadura Ikeda). The varieties were specifically selected for their range of divergent traits, including plant growth habit, fruit size and shape and pungency: Habanero peppers are popular in the Yucatán peninsula of Mexico and some islands of the Caribbean and their fruits are characterized by strong pungency and fruity aroma. Cultivar Biquinho is a sweet pepper extremely popular in Brazil, where it is sold fresh or pickled as snack. Jalapeños are signature hot peppers which account for 30% of Mexico’s hot pepper production (Sandoval-Castro et al., 2017), whereas Cascadura Ikeda is a traditional sweet bell pepper bred in Brazil. We analysed vegetative and reproductive growth traits, fruit size and shape parameters, and performed the metabolic profiling of fruit placentas and pericarps from full diallel (reciprocal) crosses between cultivars of the same or different species. We found that F_1_ hybrids show agronomic traits whose phenotypic values in many cases are heterotic with respect their parents, and with a marked reciprocal effect. Lastly, we found a strong influence of the pollen genotype on fruit size and shape of hybrids. We discuss our findings in the context of the use of intraspecific and wide crosses for breeding new varieties of *Capsicum* with novel fruit traits and increased levels of pungency.

## Results

### Hybridization impacts vegetative growth, yield and drives distinct patterns of correlation between phenotypic traits

The parental genotypes (see Supplemental Table S1 for details) show contrasting growth habit: the cultivars of *C. chinense* (HAB, BIQ) display a predominantly horizontal canopy architecture, with more vigorous stems and smaller fruits (Fig. 1A). The cultivars of *C. annuum* (JAL, IKE) have instead a more upright canopy architecture and larger fruits (Fig. 1A). Intraspecific hybridization in *C. chinense* resulted in shorter and wider plants, whereas in *C. annuum* it produced plants as tall as the JAL parent, but with increased lateral growth (Fig. 1B). Interspecific hybrids showed increased vegetative development, with greater vertical (height) and horizontal (diameter) growth (Fig. 1C), greater leaf area, resulting in greater dry vegetative biomass but lower yield at the time of harvest (Table 1). We found higher leaf area and plant diameter in both intra- and interspecific hybrids (Table 1). Biomass partitioning was significantly different between parents and hybrids: Parents of both species showed high allocation to reproductive parts, whereas interspecific hybrids had increased vegetative biomass (Supplemental Fig. S1). All interspecific hybrids showed an increase in vegetative growth, mainly in stems and roots but also leaves, with severely reduced allocation to fruits (Supplemental Fig. S1). We found a marked reciprocal effect for vegetative growth in the JAL × IKE and HAB × JAL hybrids, which was increased compared to the IKE × JAL and JAL × HAB reciprocal hybrids (Supplemental Table S2).

**Figure 1.**
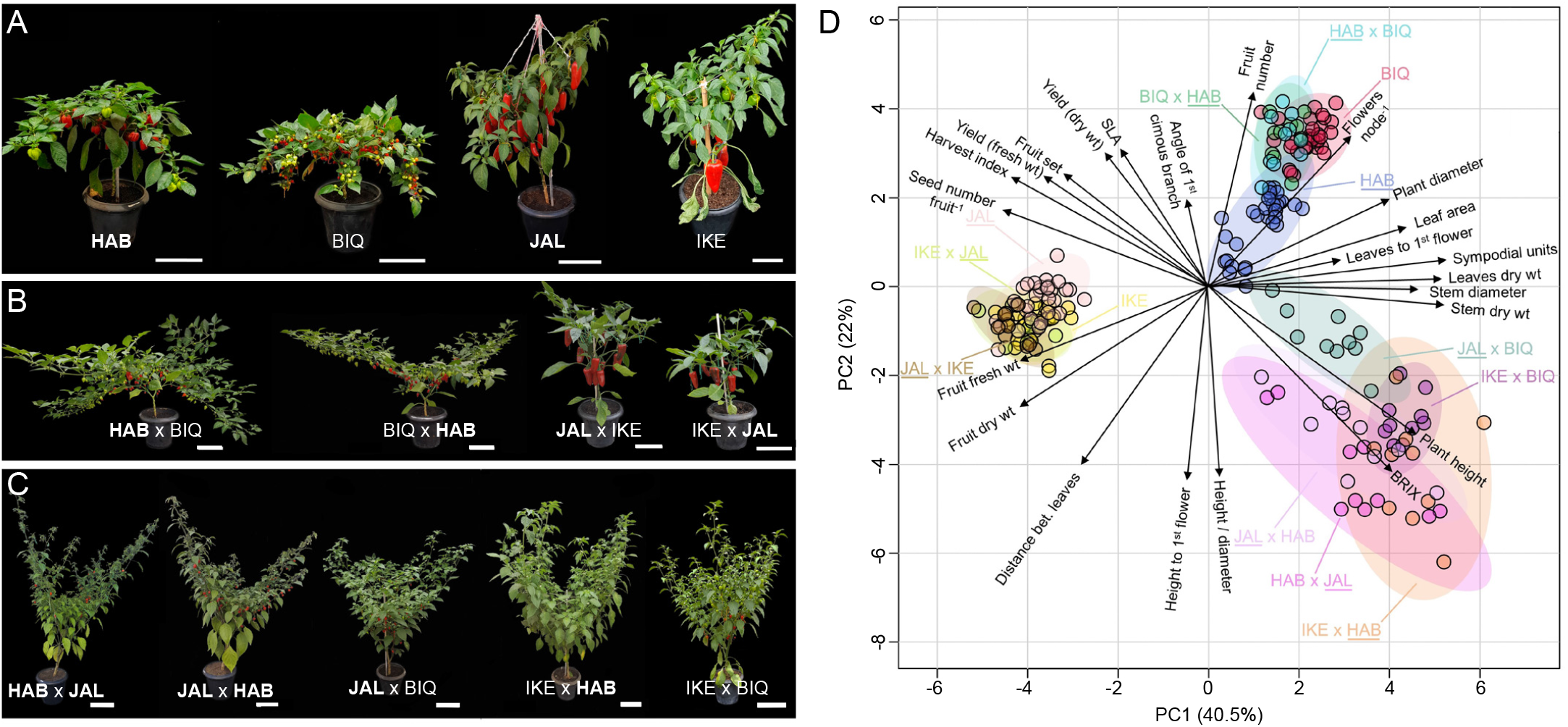
Representative *Capsicum* plants and phenotypic analysis of their growth and agronomic traits. A, representative plants of the parental genotypes: *Capsicum chinense* cv. Habanero (HAB) and cv. Biquinho (BIQ), *C. annuum* cv. Jalapeño (JAL) and cv. Ikeda (IKE). B, intraspecific hybrids; C, interspecific hybrids. Scale bars = 15 cm. The genotypes with pungent fruits are shown in **boldface**. D, PCA biplot of 13 *Capsicum* genotypes (parents, intraspecific and interspecific hybrids), based on the variance of the 28 traits listed in Supplemental Table S3, explained by two PC axes (PC1 and PC2). The two first components explain 40.5% and 22% of the variance, respectively. Different colours represent the different genotypes, pungent genotypes are <u>underlined</u>. Aligned vectors indicate positive and perpendicular vector negative correlation between traits. The numbers represent each individual sample, listed in the Supplemental Data files.

**Table 1.** Agronomic traits in intra- and interspecific hybrids of *Capsicum chinense* cv. Habanero (HAB) and cv. Biquinho (BIQ); and *C. annuum* cv. Jalapeño (JAL), and cv. Cascadura Ikeda (IKE) determined 180 days after sowing. The pungent genotypes are shown in **boldface**. Values are means ± s.e.m. (n=10 plants). Different letters within each column indicate significant differences by the Scott-Knot test at 5%.

| Genotypes | Plant height (cm) | Plant diameter (cm) | Leaf area (m<sup>2</sup>) | Yield (g) |
| --- | --- | --- | --- | --- |
| *Parents* |  |  |  |  |
| **HAB** | 59.70 ± 2.13 **f** | 94.7 ± 2.09 **d** | 0.664 ± 0.03 **d** | 862.00 ± 65.75 **c** |
| BIQ | 65.56 ± 1.65 **f** | 104.22 ± 4.80 **d** | 0.710 ± 0.03 **d** | 810.11 ± 31.93 **c** |
| **JAL** | 66.80 ± 2.67 **f** | 71.40 ± 2.53 **e** | 0.368 ± 0.02 **f** | 1243.47 ± 78.09 **b** |
| IKE | 80.89 ± 2.58 **e** | 61.11 ± 2.05 **e** | 0.576 ± 0.02 **e** | 2097.83 ± 75.58 **a** |
| *Intraspecific F<sub>1</sub>* |  |  |  |  |
| **HAB** × BIQ | 44.75 ± 4.64 **g** | 165.90 ± 6.5 **a** | 0.946 ± 0.03 **c** | 927.32 ± 57.16 **c** |
| BIQ × **HAB** | 44.65 ± 3.37 **g** | 154.5 ± 6.15 **b** | 0.867 ± 0.04 **c** | 943.78 ± 48.74 **c** |
| **JAL** × IKE | 77.60 ± 4.49 **e** | 80.70 ± 1.83 **e** | 0.362 ± 0.03 **f** | 596.99 ± 56.85 **d** |
| IKE × **JAL** | 71.10 ± 4.39 **f** | 68.3 ± 3.02 **e** | 0.364 ± 0.02 **f** | 439.98 ± 62.13 **e** |
| *Interspecific F<sub>1</sub>* |  |  |  |  |
| **HAB** × **JAL** | 135.80 ± 6.38 **c** | 122.10 ± 4.5 **c** | 0.820 ± 0.06 **c** | 197.05 ± 56.99 **f** |
| **JAL** × **HAB** | 106.90 ± 5.01 **d** | 100.8 ± 5.66 **d** | 0.754 ± 0.04 **d** | 263.51 ± 55.74 **e** |
| **JAL** × BIQ | 134.0 ± 6.28 **c** | 133.6 ± 5.04 **c** | 0.842 ± 0.09 **c** | 419.83 ± 98.72 **e** |
| IKE × **HAB** | 161.47 ± 7.51 **b** | 176.13 ± 12.10 **a** | 1.407 ± 0.11 **a** | 103.09 ± 34.59 **f** |
| IKE × BIQ | 174.430 ± 2.47 **a** | 170.7 ± 5.67 **a** | 1.232 ± 0.03 **b** | 280.51 ± 49.26 **f** |

Our initial analysis of 28 agronomic and growth parameters suggested that the main impact on yield in hybrids, particularly interspecific ones, was produced by increased vegetative growth (Supplemental Fig. S1). We next performed a principal component analysis (PCA) to reduce the dimensionality of the dataset and identify other potential traits that contribute most to the differenced observed between parents and hybrids (Fig. 1D). The genotypes were divided in the 2D scores plot into three groups: *C. chinense* cultivars and their hybrids, *C. annuum* cultivars and their hybrids and interspecific hybrids (Fig. 1D). More than 80% of the variance was captured by the first four PCA axes, so we retained PC1 to PC4 for analysis (Supplemental Table S3).

PC1 explained 40.5% of the total variability between traits and genotypes and was associated mainly with vegetative traits: dry weight of the plant parts, number of sympodial units, plant width and leaf area. Confirming our initial observations, the most significant trait for PC1 was stem dry weight. All the *C. annuum* genotypes (parents and intraspecific hybrids) grouped at the lower end of PC1 and the interspecific hybrids at the higher end, as these were the genotypes with higher levels of vegetative growth, mainly driven by increased stem weight. The PC1 divided plant height and yield, where the taller genotypes were the least productive. PC2 accounted for 22% of the total variability among traits and was mostly related to plant height and fruit total soluble solids (Brix). Interspecific hybrids grouped at the higher end of PC2, while *C. chinense* genotypes (parents and intraspecific hybrids) grouped at the lower end. Hierarchical clustering analysis of the 28 traits confirmed the strong association between yield, harvest index, fruit set and number of seeds per fruit (Supplemental Fig. S2).

### Hybridization conditions seed and fruit development in a genotype-specific manner

All crosses set seeds successfully, and seed size was higher in *C. annuum* than in *C. chinense* cultivars and their respective intraspecific hybrids (Fig. 2A). Although seed weight was strongly determined by the female parent genotype, some paternal effect was observed for interspecific crosses, *e*.*g. C. annuum* pollen on *C. chinense* pistils decreased seed size, HAB pollen on BIQ pistils increased seed size and *C. chinense* pollen increased seed size in JAL pistils (Fig. 2). For interspecific hybrids, using *C. annuum* as pistillate parent and *C. chinense* as pollen donor led to higher seed weight and the reciprocal cross to lower seed weight. Germination was higher in *C. annuum* and lower in *C. chinense* intraspecific hybrids and severely compromised in interspecific hybrids (Fig. 2B). We observed strong unilateral incompatibility in seed germination: using *C. chinense* as pistillate parent and *C. annuum* as pollen donor had a more negative effect than the reciprocal cross. Three of the interspecific hybrid seeds failed to germinate (BIQ × JAL, HAB × IKE and BIQ × IKE), so these genotypes are absent from all subsequent analyses.

**Figure 2.**
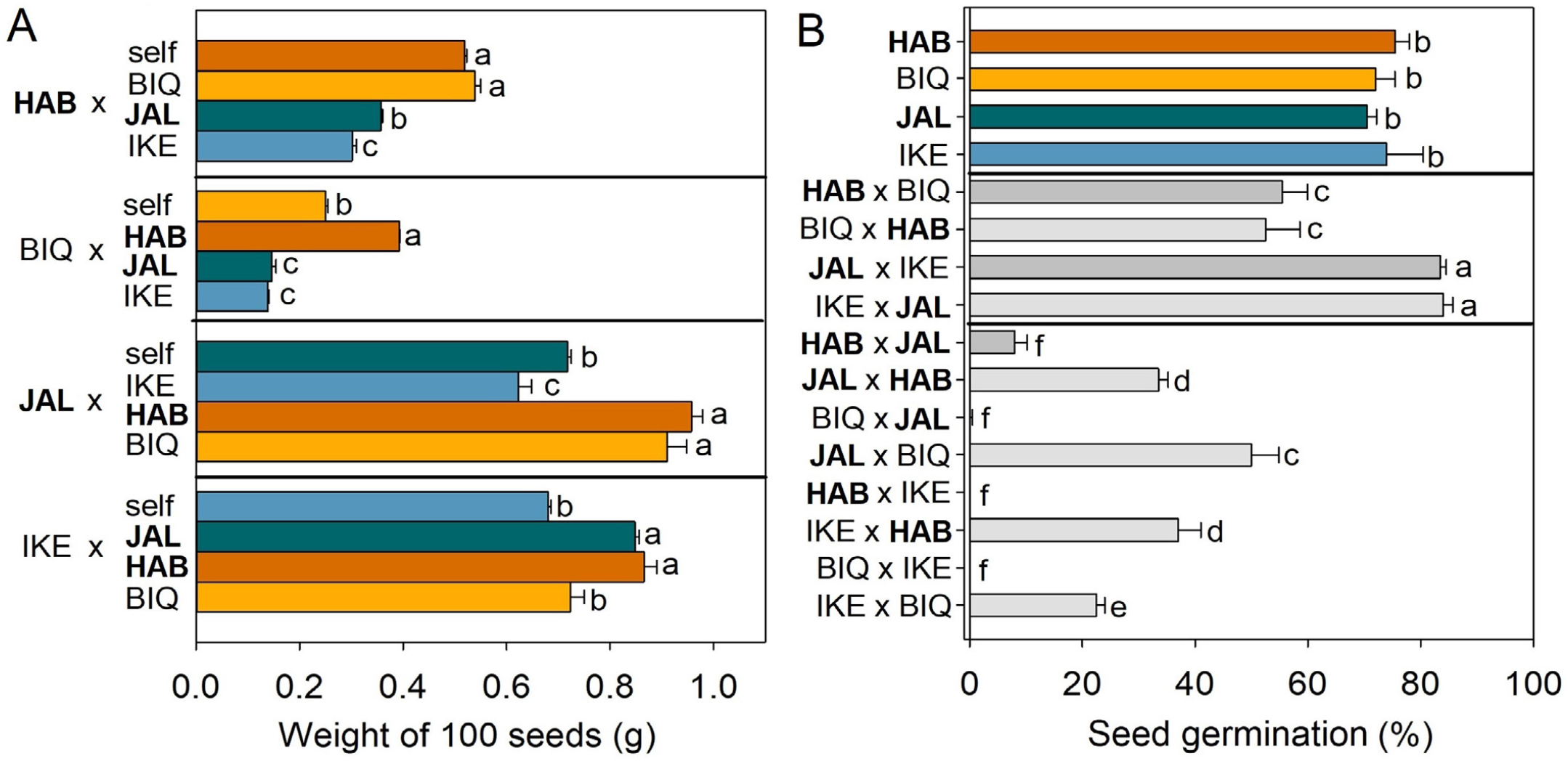
Seed traits in *Capsicum* cultivars and their hybrids. A, Seed weight and B, germination in *Capsicum chinense* cv. Habanero (HAB), cv. Biquinho (BIQ), *C. annuum* cv. Jalapeño (JAL), and cv. Ikeda (IKE) and their intra- and interspecific hybrids. The first genotype in each cross represents the pistillate parent. The pungent genotypes are shown in **boldface**. Different letters differ by Scott Knott’s 5% test.

Seed weight did not appear to have a strong effect on germination: for instance, in spite of large weight differences between parental genotypes seeds, there was no difference in germination between them (Fig. 2). By contrast, seed weight was strongly correlated to initial vegetative growth parameters (dry matter, plant height and diameter, leaf area) (Supplemental Fig. S3). Given the low fertility of interspecific F_1_ hybrids, we conducted controlled pollination with either pollen from the hybrid itself or of either parent. Remarkably, the pollen genotype had a strong effect on fruit size and shape (Supplemental Fig. S4A). Pollination with parents was an effective method of restoring fertility, as fruit set and number of seeds per fruit were increased (Supplemental Fig. S4B).

### Capsicum hybrids display heterosis and reciprocal effects for agronomic traits

Heterosis, or the superiority of hybrids to parents can be expressed as either relative mid-parent heterosis (rMPH) or relative best-parent heterosis (rBPH). We calculated these values for the intra- and interspecific hybrids and found significant rMPH and rBPH for many traits but also hybrid depression, depending on the trait and the type of cross (Table 2). The *C. chinense* intraspecific hybrids showed reduced plant height, with no difference in this trait for *C. annuum*, whereas all the interspecific hybrids were taller. Plant diameter was also increased in all hybrids. Total vegetative dry mass was decreased in intra-but increased in interspecific hybrids. Fruit set was generally decreased in all hybrids with varying intensity, with the result that fruit number was also reduced. showed reduced plant height and increased diameter. Except for *C. chinense* hybrids, which showed a slight increase, all hybrids suffered yield penalty, with a particularly severe reduction in individual fruit weight (Table 2). Total soluble solids content in the fruits (Brix), however, was increased in all hybrids except the *C. chinense* intraspecific ones. We observed reciprocal effects in many agronomic traits for the *C. annuum* but not for the *C. chinense* hybrids (Supplemental Table S3). The remarkable changes in fruit size and shape inspired us to conduct a more in-depth analysis of these agronomically important traits.

**Table 2.** Relative mid-parental heterosis (rMPH) and best-parental heterosis (rBPH) for twelve traits in two intraspecific and four interspecific hybrids of *Capsicum*. Parents: *C. chinense* cv. Habanero (HAB), *C. chinense* cv. Biquinho (BIQ), *C. annuum* cv. Jalapeño (JAL) and *C. annuum* cv. Cascadura Ikeda (IKE). The pungent genotypes are shown in **boldface**. The significance test for the variables rMPH and rBPH was performed based on the absolute values of MPH and BPH. *, ** indicate significant values at *p* = 0.05 and 0.01, respectively by the Scheffe test.

|  | Hybrids |  |  |  |  |  |
| --- | --- | --- | --- | --- | --- | --- |
|  | Intraspecific F<sub>1</sub> |  | Interspecific F<sub>1</sub> |  |  |  |
|  | HAB × BIQ | JAL × IKE | HAB × JAL | JAL × BIQ | IKE × HAB | IKE × BIQ |
|  | Plant height |  |  |  |  |  |
| rMPH | -28.63 ** | 0.68 | 91.86 ** | 102.48 ** | 129.71 ** | 138.04 ** |
| rBPH | -31.81 ** | -8.08 | 81.66 ** | 100.6 ** | 99.62 ** | 115.48 ** |
|  | Plant diameter |  |  |  |  |  |
| rMPH | 61.07 ** | 12.44 * | 34.2 ** | 52.14 ** | 126.08 ** | 106.49 ** |
| rBPH | 53.71 ** | 4.34 | 17.69 ** | 28.19 ** | 85.99 ** | 63.78 ** |
|  | Dry vegetative biomass |  |  |  |  |  |
| rMPH | -23.01 ** | -41.50** | 301.12 ** | 75.28 ** | 93.36 ** | 332.77 ** |
| rBPH | -24.38 ** | -49.55 ** | 186.50 ** | 23.85 ** | 53.83 ** | 239.75 ** |
|  | Fruit set |  |  |  |  |  |
| rMPH | -2.23 | -14.67 | -70.73 ** | -54.92 ** | -99.47 ** | -83.71 ** |
| rBPH | -27.99 * | -32.73 ** | -78.04 ** | -56.16 ** | -99.60 ** | -87.42 ** |
|  | Number of fruits per plant |  |  |  |  |  |
| rMPH | -5.09 | -74.68 ** | -22.94 | -51.43 ** | -4.11 | -55.29 ** |
| rBPH | -45.01 ** | -75.03 ** | -41.8 ** | -73.75 ** | -28.26 | -75.88 ** |
|  | Yield |  |  |  |  |  |
| rMPH | 11.90 | -68.97 ** | -78.13 ** | -59.11 ** | -93.03 ** | -80.71 ** |
| rBPH | 8.53 | -75.28 ** | -81.48 ** | -66.24 ** | -95.09 ** | -86.63 ** |
|  | Fresh weight per fruit |  |  |  |  |  |
| rMPH | -38.97 ** | -22.90 * | -78.00 ** | -86.38 ** | -92.14 ** | -92.72 ** |
| rBPH | -63.42 ** | -46.55 ** | -85.05 ** | -92.70 ** | -95.53 ** | -96.26 ** |
|  | Brix ° |  |  |  |  |  |
| rMPH | -7.21 ** | 3.86 ** | 31.48 ** | 34.14 ** | 16.02 ** | 31.26 ** |
| rBPH | -17.18** | 1.56 | 19.89 ** | 30.96 ** | 8.01 | 25.3 ** |

### Fruit shape shows non-additive inheritance

Peppers are a genus with considerable fruit morphological variation, that is strongly associated with consumer preferences (Paran and van der Knaap, 2007). Thus, we next turned our attention to the effects of hybridization on fruit size and shape. The parental genotypes are highly contrasting in both traits (Fig. 3A). Individual fruit size of intraspecific hybrids was intermediate between that of the parental cultivars, whereas it was highly reduced in interspecific hybrids (Fig. 3B). Remarkably, the intraspecific hybrids resembled more closely one of the parents, regardless of the direction of the cross: BIQ in *C. chinense* hybrids and IKE in *C. annuum* (Fig. 3A). Given the multidimensional nature of fruit shape, we used TomatoAnalyzer (Rodríguez et al., 2010) to process scanned images of fruit sections and to provide an objective, quantitative assessment based on 32 fruit morphology traits. We conducted a PCA on the traits and determined four main axes that could explain 91.3% of the variability in fruit size and shape (Fig. 3C).

**Figure 3.**
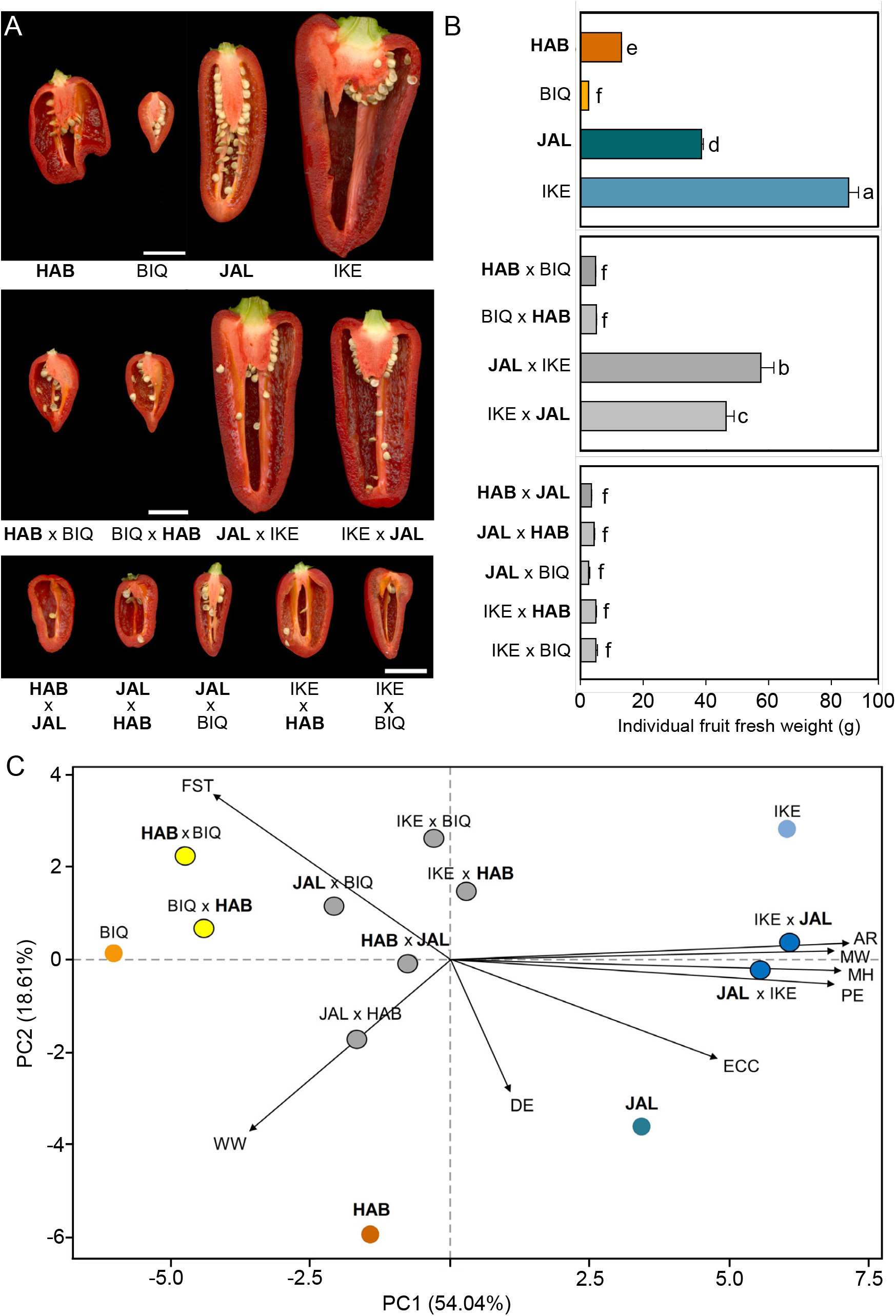
Fruit size and shape analysis in *Capsicum chinense* cv. Habanero (HAB) and cv. Biquinho (BIQ) and *C. annuum* cv. Jalapeño (JAL) and cv. Cascadura Ikeda (IKE), and their intra- and interspecific F_1_ hybrids. A, Representative fruit cross-sections of the parental lines (top), intraspecific hybrids (middle) and interspecific hybrids (bottom). The pungent genotypes are shown in **boldface**. Scale bars 2 = cm. B, individual fruit fresh weight (g). Bars are means ± s.e.m. (n=10). Letters show significant differences by ANOVA followed by Scot Knott’s 5% test. C, PCA biplot of 13 *Capsicum* genotypes (parents, intraspecific and interspecific hybrids) based on the variance in 32 fruit biometric parameters, explained by two PC axes (PC1 and PC2). The two components explained 54.04% and 18.61% of the variance, respectively. Parental genotypes are represented by circles of different colours, intraspecific *C. annuum* hybrids by blue with black outline, *C. chinense* by red with black outline and interspecific hybrids in gray. The pungent genotypes are shown in **boldface**.

PC1 accounted for 54.4% of the variability and was significantly associated with traits related to fruit size: cross-sectional fruit length and perimeter. Thus, the parental genotypes were divided along the PC1 according to fruit size (Fig. 3C). PC2 accounted for 18.6% of the fruit phenotype variability and was mostly associated with fruit shape (asymmetry, homogeneity and blockiness) (Supplemental Table S4). With its characteristic blocky, irregular fruit, *C. chinense* cv. HAB was located low along the PC2 axis and separated from all other genotypes. PC3 accounted for 10.7% of the variability in fruit shape and was associated with internal eccentricity and homogeneity traits and PC4 accounted for 7.6% of the variability with influence from pericarp traits. Our analysis also showed that the *C. chinense* intraspecific hybrids clustered very close to the BIQ cultivar, as suggested by their similarity in fruit phenotype (Fig. 3C). All the interspecific hybrid fruits were also located closer to BIQ than any other parental genotype due to their reduced size and their generally more triangular shape.

### Heterosis for pungency and primary metabolites in F1 hybrids is characterized by patterns of non-additive accumulation

Capsicum fruits are noted for their pungency (‘heat’), which is conferred by a class of metabolites collectively known as capsaicinoids, which accumulate in the placenta of pungent fruits. Since capsaicinoids are synthesized from amino acidic precursors, we carried out a detailed metabolic profiling of both polar and lipophilic primary metabolites in parental genotypes and their hybrids (Supplemental Fig. S5). As expected, the placental tissues of the pungent parents from both *C. chinense* (HAB) and C. *annuum* (JAL) showed high accumulation of the main capsaicinoids, in contrast to the sweet pepper BIQ and IKE (Supplemental Fig. S4). The most abundant capsaicinoids were capsaicin (C_18_H_27_NO_3_, [M+H]+=306.206356, dihydrocapsaicin (C_18_H_29_NO_3_, [M+H]+=308.222006) and nordihydrocapsaicin (C_17_H_27_NO_3_ [M+H]+=294.206356), which accounted for over 90% of total capsaicinoid content in all genotypes (Fig. 4). All hybrids were pungent, regardless of their parental genotypes and, except for the intraspecific hybrids of *C. annuum*, all combinations displayed significant degrees of both rMPH and rBPH for all of the main capsaicinoids (Table 3). Rather interestingly, the IKE × BIQ F_1_, which derives from the cross of two non-pungent parents, shows a high accumulation of capsaicinoids.

**Figure 4.**
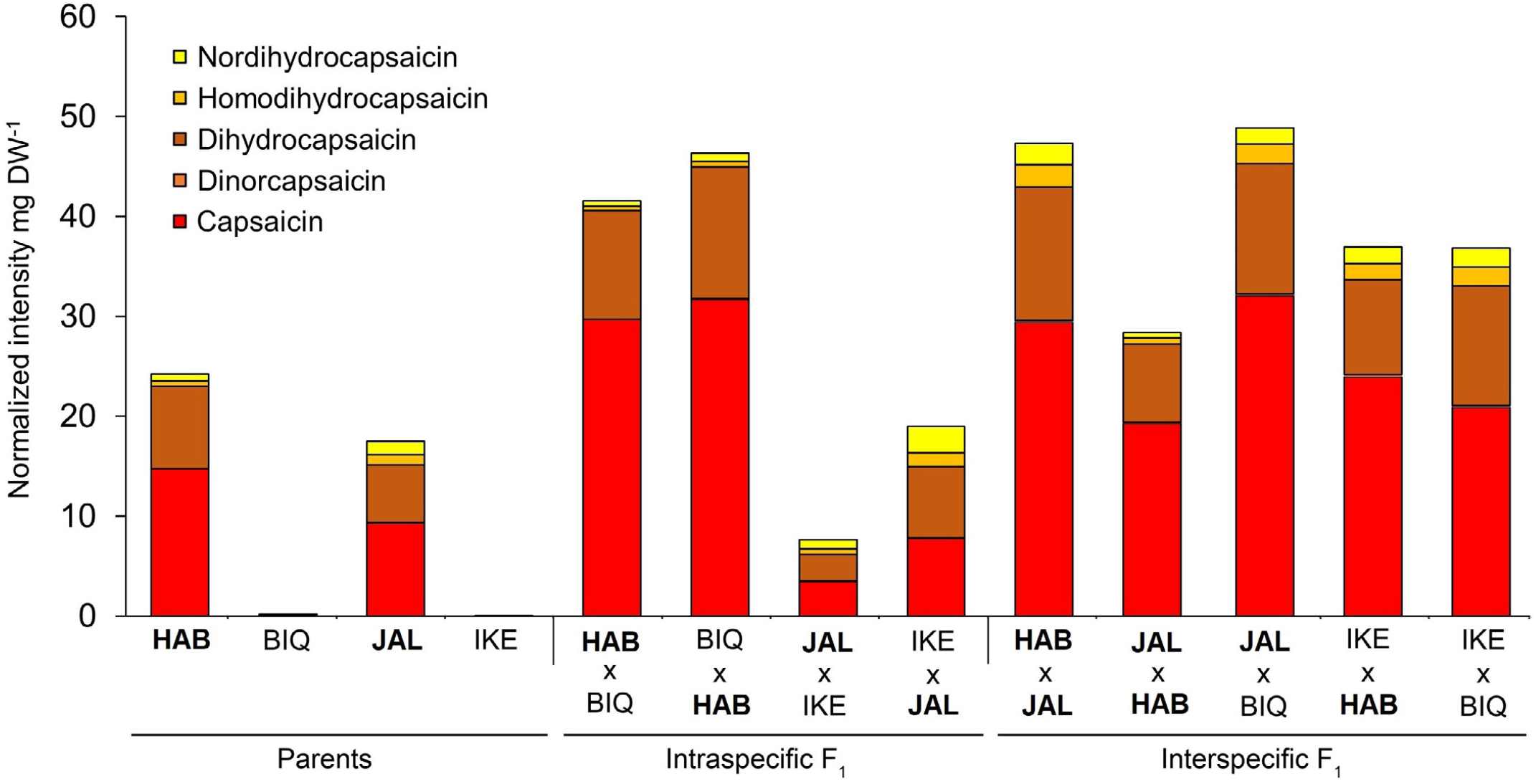
Relative capsaicinoid content in the placenta of *Capsicum chinense* cv. Habanero (HAB), cv. Biquinho (BIQ), and *C. annuum* cv. Jalapeño (JAL) and cv. Cascadura Ikeda (IKE) and their intra- and interspecific hybrids. The pungent genotypes are shown in **boldface.** Each column represents the mean value per genotype (n=7).

**Table 3.** Relative mid-parental heterosis (rMPH) and best-parental heterosis (rBPH) for the accumulation of the three main capsaicinoids (capsaicin, dihydrocapsaicin and nordydrocapsacin) and for the total capsaicinoids in the placenta of fully ripe fruits in intraspecific and interspecific hybrids of *Capsicum*. Parents: *C. chinense* cv. Habanero (HAB), *C. chinense* cv. Biquinho (BIQ), *C. annuum* cv. Jalapeño (JAL) and *C. annuum* cv. Cascadura Ikeda (IKE). The pungent genotypes are shown in **boldface**.

| Hybrids |
| --- |

| Genotype | Intraspecific F<sub>1</sub> |  |  |  | Interspecific F<sub>1</sub> |  |  |  |  |
| --- | --- | --- | --- | --- | --- | --- | --- | --- | --- |
|  | **HAB** ×<br>BIQ | BIQ ×<br>**HAB** | **JAL** ×<br>IKE | IKE ×<br>**JAL** | **HAB** ×<br>**JAL** | **JAL** ×<br>**HAB** | **JAL** ×<br>BIQ | IKE ×<br>**HAB** | IKE ×<br>BIQ |
| Capsaicin |  |  |  |  |  |  |  |  |  |
| rMPH | 300.70<br>** | 328.46<br>** | -34.7 | 54.59 | 132.16 ** | 168.11 ** | 347.05 ** | 184.16 ** | 39493.63<br>** |
| rBPH | 101.65<br>** | 115.63<br>** | -67.35<br>** | -22.69 | 99.90 ** | 130.85 ** | 125.54 ** | 42.10 ** | 20027.80<br>** |
| Dihydrocapsaicin |  |  |  |  |  |  |  |  |  |
| rMPH | 161.43<br>** | 217.84<br>** | -31.96 | 127.22<br>** | 79.89 ** | 75.87 ** | 186.84 ** | 189.16 ** | 76568.97<br>** |
| rBPH | 31.03<br>** | 59.30 ** | -65.98<br>** | 13.62 | 61.71 ** | 58.10 ** | 43.85 ** | 44.59 * | 39402.03<br>** |
| Nordihydrocapsaicin |  |  |  |  |  |  |  |  |  |
| rMPH | 58.92<br>** | 131.32<br>** | -8.59 | 307.87<br>** | 94.36 ** | 47.29 * | 117.87 ** | 453.92 ** | 46169.57<br>** |
| rBPH | -20.28<br>** | 16.04 * | -54.29<br>** | 103.94<br>** | 42.15 ** | 7.73 | 9.1 | 176.96 ** | 23034.78<br>** |
| Total Capsaicinoids |  |  |  |  |  |  |  |  |  |
| rMPH | 2745.10<br>** | 284.14<br>** | -31.64 | 100.57<br>** | 111.88 ** | 129.49 ** | 272.43 ** | 193.87 ** | 45860.17<br>** |
| rBPH | 1322.81<br>** | 93.03 ** | -65.82<br>** | 0.300 | 89.74 ** | 105.52 ** | 87.39 ** | 46.95 ** | 23034.78<br>** |

We then mapped the metabolite rBPH values onto the main pathways of primary metabolism and found that a few general trends emerged (Fig. 5 for the placenta and Supplemental Fig. S6 for the pericarp). We observed marked reciprocal effects on the level of several metabolites (mainly amino acids, e.g., serine, valine, aspartate, GABA and pyroglutamic acid) in the placenta of the fruits derived from the intraspecific *C. chinense* crosses (i.e., HAB × BIQ and its reciprocal). This reversion of the heterotic effect was not observed in the other intraspecific cross (the one derived from crossing JAL and IKE, both cultivars of *C. annuum*), where most of the primary metabolites, including amino acids, show the same sign of rBPH irrespective of whether JAL or IKE was used as the female parent (Fig. 5A). Primary metabolites from the interspecific crosses (Fig. 5B) showed instead a more uniform pattern of hybrid depression (negative rBPH), except for trehalose and asparagine, which displayed considerable rBPH in the interspecific hybrids where IKE was used as female (Fig. 5B). In the pericarp (Supplemental Fig. S6), strong rBPH was detected for serine (along with its probable precursor, glycerate, given that serine biosynthesis in non-photosynthetic tissues mainly occurs through the phosphorylated pathway (Galili et al., 2016), as well as for other amino acids derived from TCA intermediates (lysine, aspartate, asparagine and pyroglutamic acid).

**Figure 5.**
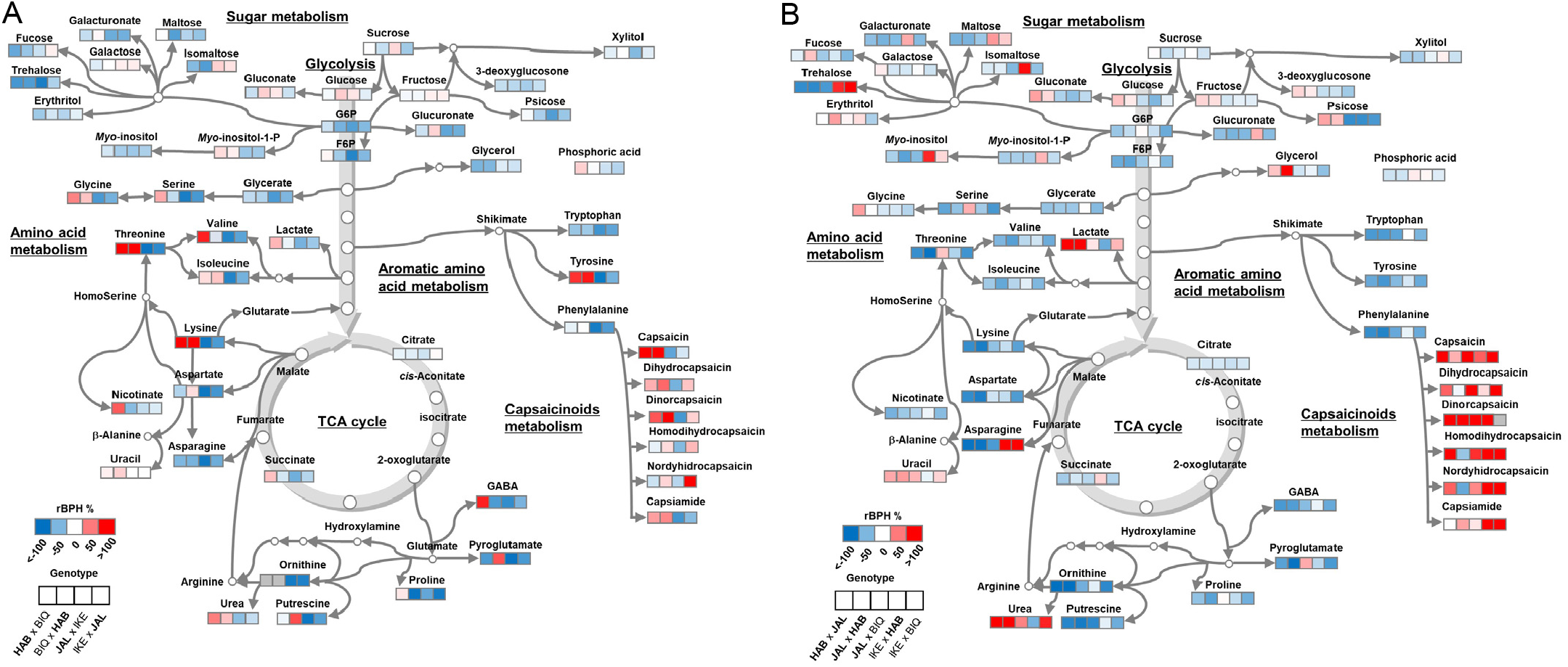
Heat map showing heterosis for primary metabolites and capsaicinoids in the fruit placenta of *Capsicum chinense* cv. Habanero (HAB), cv. Biquinho (BIQ), and *C. annuum* cv. Jalapeño (JAL) and cv. Cascadura Ikeda (IKE) hybrids. A, intraspecific and B, interspecific hybrids. The pungent genotypes are shown in **boldface**. Each square represents a genotype (n=7). The color key indicates the relative best parent heterosis (BPH) value (red = higher, blue = lower).

## Discussion

### *The impact of hybridisation on vegetative and agronomic traits in* Capsicum

The most commonly produced and consumed peppers in the world belong to the *C. annuum* var. *annuum* (here *C. annuum* for brevity), whereas most sweet peppers are varieties of *C. annuum*, some of the most pungent varieties belong to *C. chinense* (Jang et al., 2021). *C. chinense* originates in the Amazon basin and is therefore better adapted to hot and humid conditions (Pickersgill, 2007), so accessions of this species are valuable sources of multiple pathogen resistance genes (Di Dato et al., 2015). Interspecific hybridisation would be a suitable avenue to combine the favourable traits found in different species of the genus. Hybrid breeding is the simplest way of combining dominantly inherited traits. Although considerable effort has been expended to explore the agronomic consequences of intraspecific hybridisation in *C. annuum* (Marame et al., 2009; Singh et al., 2014; Tripodi et al., 2020), little is known concerning the effects of hybridisation between species. Here, we found that clearly divergent traits have been favoured in the cultivars of each species selected for this study, the most remarkable being bigger fruits of *C. annuum* and more numerous fruits of *C. chinense*. The intraspecific hybrids of each species show additive effect for fruit weight in the *C. annuum* hybrids and fruit number in the *C. chinense* ones. Our results show that this directional selection had an impact on plant vegetative traits, as evidenced by the strong negative correlation between fruit number and plant height and a positive one with plant diameter. By contrast, large fruits correlate negatively with plant and stem diameter and leaf area.

High yield was maintained in *C. chinense* hybrids but not in *C. annuum* ones and was severely compromised in interspecific hybrids. This suggests that the genetic control underlying the traits that control yield (fruit size, fruit number and fruit set) is recessive and, therefore, hybrid breeding is amenable only between cultivars where the agronomic basis for yield is similar (*i*.*e*. either fruit size or number). We also found a strong negative correlation between seed number and total soluble solids (Brix). Reducing seed number and increasing Brix are both desirable agronomic traits; the energetic cost of seed production could negatively impact fruit total solids (sugars and organic acids). High Brix is also negatively correlated with fruit size, as in tomato (Bernacchi et al., 1998).

Enhanced seed germination is a key agronomic trait in cultivated species, as opposed to their highly dormant wild ancestors (Soltani et al., 2021). When *C. annuum* was used as the female parent, all crossing combinations with *C. chinense* yielded viable hybrids. In the reciprocal crosses, when cultivars of *C. chinense* were instead used as the female parent, the only fertile hybrid we recovered was the combination HAB × JAL. This may suggest a general weakness of the maternal tissues of *C. chinense* - especially of cultivar BIQ *-* to provide nutrients to the embryo during the early developmental phases (Meyer et al., 2012). The size and viability of seeds can be affected by many different factors: the concurrent accumulation of capsaicinoids in the placenta (the tissue upon which seeds set and remain attached (Barchenger and Bosland, 2016)), but also by many other maternal effects (*e*.*g*. transport processes to the developing seeds), depending on the species. Theory predicts that seed size is determined by the conflict between the paternal and maternal genotypes (Pires and Grossniklaus, 2014). This would therefore result in a trade-off between seed size and number. Producing viable hybrids continues to be a major challenge for hybrid pepper breeding and further work needs to be conducted to understand the genetic and physiological basis of post-zygotic hybridization barriers (Lafon-Placette and Köhler, 2016).

### *Heterosis and reciprocal effects in* Capsicum *hybrids*

In peppers, heterosis is most likely to be observed as an increase in vegetative mass. Here, we found increased vegetative growth in both intra- and interspecific hybrids, however the effect was more dramatic in the latter, as it resulted in a strong yield penalty. Biomass allocation to reproductive tissues is a hallmark of crop domestication and breeding (Hufford et al., 2019). Our results highlight the diverging genetic bases of breeding for high yield in *C. annuum* and *C. chinense*. Pepper breeders have used wide crosses and introgression to generate high yielding varieties (Srivastava and Mangal, 2019). As a trade-off, these crops have less allocation of biomass to stems and roots, and increased susceptibility to diseases and pests. It is an open question why this was not observed in the crosses reported here. Interspecific hybridization would be a way to introduce vigorous characteristics such as stem and root or even to gain an understanding of related mechanisms and genes related to greater vegetative vigor. One of our interspecific crosses (IKE × BIQ) produced a vigorous root system (>25% of total biomass allocation), which could be relevant within the context of breeding improved rootstocks for grafting production systems (Kyriacou et al., 2017).

The asymmetry between the phenotypic value of certain traits (Supplemental Table S3) of reciprocal crosses is attributed to reciprocal effects, which could be related to cytoplasmic inheritance or nucleus-cytoplasm incompatibility mechanisms (Joseph et al., 2013). We found more marked reciprocal effects for *C. annuum* than for *C. chinense*, particularly for vegetative biomass, fresh weight per fruit, number of seeds per fruit and Brix. Unilateral incompatibility is an extended phenomenon in *Capsicum* peppers (Onus and Pickersgill, 2004), however *C. annuum* and *C. chinense* are generally considered to be fully compatible. Here, we have shown that using *C. annuum* as a pistillate (female) parent leads to successful production of interspecific hybrids with *C. chinense* but the reciprocal type of cross is hampered by strong incompatibility. Further work is required to understand the underlying biological mechanisms and aid hybrid breeding in *Capsicum*.

We found considerable heterosis specifically for capsaicinoids accumulation in the placenta of both intra- and interspecific hybrids, including those derived from non-pungent parents (BIQ and IKE). This behavior of non-additive heterotic activation in the F_1_ may be due to the complementation of non-functional alleles at distinct loci, with the restoration of a fully functional capsaicinoid pathway in the hybrids. By contrast, the pericarp metabolites of interspecific hybrids showed mid- to strong hybrid depression, except for several amino acids from the HAB × JAL combination (glycine, threonine, branched-chain and aromatic amino acids, lysine, aspartate, asparagine, GABA, pyroglutamic acid and GABA). Recent work on *Brassica juncea* showed that primary and secondary metabolites display both additive and non-additive inheritance, depending on tissue type (buds vs. leaves) and developmental stage (Bajpai et al. 2019). Here, most primary metabolites, whether from placenta or pericarp, generally showed non-additive accumulation (i.e., their levels being significantly lower or higher with respect to the best parent), with few cases of reciprocal effects detected for amino acids in the placenta of the intraspecific cross of *C. chinense*.

### The effect of hybridization on fruit size and shape

*Capsicum* is a genus recognized and appreciated by consumers for its diversity in fruit shape and color (Scossa et al., 2019). To a large extent, the diversification is related to the geographic expansion and selection for culinary and aesthetic traits within the genus (Tripodi et al., 2021). Cultivars of *C. chinense* show high diversity in fruit formats and growth parameters (Rosado-Souza et al., 2015; Bianchi et al., 2020). The genetic basis of fruit size and shape is hitherto unknown in peppers, although an effort is underway to fill this gap in the knowledge (Colonna et al., 2019; Nimmakayala et al., 2021). Fruit shape is a multidimensional trait, so we broke it down in its individual components using the TomatoAnalyzer resource (Rodríguez et al., 2010). We then conducted a principal component analysis (PCA) and, as expected, the first principal component discriminated all parental cultivars mostly based on fruit size (cross sectional area, perimeter, width and length). By contrast, the second principal component was dominated by shape traits (blockiness, ellipsoid, asymmetry, eccentricity). Interestingly, the intraspecific hybrid fruits grouped together with the non-pungent parents, even when both were at the opposite extremes of the fruit size distribution (BIQ, smallest and IKE, biggest). This suggests that dominant genetic variance is a key component of fruit traits in non-pungent *Capsicum* varieties. Thus, our results show the feasibility of easily creating pungent fruits that retain the highly valued visual and organoleptic traits of non-pungent varieties.

Surprisingly, the pollen genotype had a strong impact on fruit size and shape. This apparently anomalous phenomenon was first described by Darwin as “the direct action of the male element on the female form” and subsequently called ‘xenia’. Xenia is widespread in cultivated species, including cereals, vegetables and fruit trees (Trueman et al., 2021). To the best of our knowledge, however, no reports exist of xenia in *Capsicum* (Liu, 2018). It is potentially a highly relevant phenomenon for plant breeding and crop production, for instance, pollen origin can increase yield in beans (Duc et al., 2001), maize (Weingartner et al., 2002) and raspberry (Żurawicz et al., 2018). The molecular mechanism underlying xenia is unknown, and it has been proposed that small mobile RNAs derived from the pollen grain could trigger a massive reprogramming of gene expression in the zygote and embryo, which could in turn influence the surrounding ovary tissue. Our scheme of backcrosses of *Capsicum* interspecific hybrids could represent a useful model system to explore this topic.

The variation in shape, size and colour of pepper fruits can act as a marker for variation in other, less evident characters, such as pungency. This phenomenon whereby combinations of characters allow fruits to be distinguished is called ‘perceptual distinctiveness’ (Boster, 1985) and has been described in peppers and other species (Pickersgill, 2007). For a new cultivar to be accepted, it must be identifiable on the basis of a suite of characters and distinguishable from those already in cultivation. Pepper diversity depends on morphometric and colorimetric trait variation, which is critical for varietal identification and breeding (Nankar et al., 2020). Our results show that the strong association of small peppers with the characteristic ‘biquinho’ (‘little beak’) shape or the large blocky bell peppers with sweetness can be broken down to create hybrid varieties with a similar shape, but that are highly pungent.

## Conclusion

Hybrid breeding has been underexploited in *Capsicum*, in large part due to a lack of knowledge about heterosis and reciprocal effects in different hybrid combinations. With the increasing challenge of unpredictable climate, exploiting the full range of natural variation for peppers and other crops will be a suitable avenue to create novel, resilient varieties. It can also pave the way for the creation of novel hybrids with altered combinations of visual and organoleptic traits, such as fruit size, shape, colour and flavour. Here, we have shown that combinations of *C. annuum* var. *annuum* and *C. chinense* commercial varieties can produce valuable new agronomic phenotypes through heterosis and reciprocal effects. Further exploration of the genetic basis of these phenomena will contribute to the knowledge-based breeding of more resilient and appealing pepper varieties.

## Material and methods

### Plant material and growth conditions

Two cultivars of *Capsicum annuum* var. *annuum* (henceforth called *C. annuum* for brevity) were cultivated: the sweet bell pepper “Cascadura Ikeda” (IKE) and the pungent “Jalapeño” (JAL), while for *C. chinense*, the pungent cultivar “Habanero” (HAB) and the sweet “Biquinho” (BIQ) were grown (Table S1). Intra- and interspecific F1 hybrids were created following a full diallel crossing scheme, by emasculating the flower buds before anthesis, and then transferring pollen of the selected male parent. The experiments were conducted in a greenhouse at the Federal University of Viçosa, between August 2018 and December 2019. For productivity analysis, each *Capsicum* hybrid and its respective parents (controls) were grown until harvest (when 70% of the fruits reached the ripe state) in 7.5 L pots containing the commercial substrate Tropostrato® and irrigated daily and fertilized with 4.5 g L^-1^ of NPK (4-14-8) and 4 g L^-1^ of dolomitic limestone. Supplemental fertilization was provided fortnightly with foliar spray. The compatibility between the parents, as well as the compatibility between the F_1_ hybrids and their respective parents was measured through crosses in all possible combinations: 25 flowers were pollinated of four different plants per genotype (100 flowers per cross). As a control, 100 flowers were tagged but self-pollinated. The fresh weight, length, diameter and number of seeds per fruit were estimated for the 12 largest fruits among all produced from each controlled cross.

### Seed germination assays

To test seed germination, we sowed 100 seeds per block, with a total of four blocks. We pre-treated the seeds with 3% sodium hypochlorite and subsequently washed them with abundant deionized water before sowing. We sowed the seeds in plastic pots containing the commercial substrate Tropstrato® and kept them in greenhouse conditions in the summer. We counted the total number of emerged plants 30 days after the sowing date and estimated the germination percentage.

### Vegetative growth and agronomic determinations

Thirty days after germination plants were transplanted to 7.5 L pots with Tropstrate commercial soil. Plants were grown to maturity and the following measurements were conducted on ten plants per genotype: plant height (height of soil base up to leaf apex), canopy diameter, stem diameter, number of leaves, leaf area, root, stem and leaves dry mass, stem diameter, number of leaves, soil base height up to the leaf apex and canopy diameter, number, time to anthesis and to first ripe fruit. At the time of harvest, productivity was determined as number and in total fresh weight of fruits per plant. The angle formed between the first ten branches was measured in addition to the total number of sympodial units per plant.

### Fertility Estimation

We measured fruit set (%), fruit size, number of seeds per fruit and seed germination rate. Fruit set was measured as the percentage of fruits developed from marked flowers at the time of anthesis. The average mass (g) of 100 F_1_ seeds was also quantified (*n* = 6) for the seeds resulting from crosses as self-pollination. The seeds were removed from fully ripe fruits and air-drying until they reached a constant weight. Germination tests were performed to verify the relationship between seed production, number, size and germination with fertility. One hundred seeds per replicate were sown 30 days after sowing, the number of emerged seedlings was counted and expressed as the percentage germination.

### Determination of fruit total soluble solids (Brix)

Six fully ripe fruits per plant were harvested, totaling 60 fruits per genotype. The ºBrix was determined through the digital bench refractometer model RTD-45 (Instrutherm, São Paulo, Brazil). The analyses were performed after calibrating the refractometer with distilled water and obtaining the zero, after which the collector part was wiped clean, then the pericarp was squeezed to extract enough liquid to cover the prismatic surface of the digital refractometer.

### Metabolic analyses of fruits

Flowers were tagged at anthesis and samples of fruit pericarp and placenta were collected at midday after 60 days. The pericarp and placenta from seven fruits per genotype were collected from different plants and used for the analyses. Samples were immediately frozen in liquid nitrogen and stored at −80 °C. The material was then freeze-dried (Scanvac, Coolsafe 55-4) and ground to a fine powder. The metabolic profile from each sample was determined following previously described procedures (Salem et al., 2017; Vijlder et al., 2018; Lapidot-Cohen et al., 2020), with modifications summarized briefly here. 10 mg of sample material was mixed with 1 mL of extraction buffer consisting of methyl-*tert*-butyl-ether and methanol (3:1), and containing an internal standard of 50 μL corticosterone (1 mg mL^-1^ in methanol), 50 μL 1,2-diheptadecanoyl-*sn*-glycero-3-phosphocholine (1 mg mL^-1^ in chloroform) and 50 μL ribitol (1 mg mL^-1^ in water). Samples were vortexed and incubated in a shaker (1000 rpm) at 4°C for 45 min. 650 μL of water:methanol (v/v 3:1) were then added to induce phase separation and the samples were centrifuged at 20000 *g* for 5 min at room temperature. We then transferred aliquots from both the upper (apolar) and lower phase (polar and semi-polar metabolites) to new tubes to be reduced to dryness in the speedvac for subsequent metabolic analyses. The dried aliquots were stored at −80°C until derivatization or resuspension prior to GC- or LC-MS analyses.

For the analysis of primary metabolites, dried aliquots from the polar liquid phase were initially derivatized with 60 μL of methoxiamine hydrochloride (30 mg mL^-1^ in pyridine) and shaken at 37°C for two hours. Sample extracts were then trimethylsylilated with 120 μL of a mix of methyl-*N*-(trimethylsylil)trifluoroacetoamide (MSTFA) containing standards of fatty acid methyl esters (FAMEs), followed by agitation at 37°C for 30 minutes. Samples were spun briefly and a volume of 140 μL from each was transferred to glass vials and 1 μl injected for gas chromatography-time of flight-mass spectrometry (GC-TOF-MS) as described previously (Osorio et al., 2012). Chromatograms and mass spectra were evaluated using ChromaTOF 1.0 (Leco, www.leco.com) and TagFinder v.4.0, respectively. Cross-referencing of mass spectra was performed with the Golm Metabolome database (Kopka et al., 2005).

For the analysis of apolar metabolites, dried aliquots from the upper lipid phase were resuspended in 400 μL of acetonitrile:2-propanol 7:3 (v/v) and 140 ul were then transferred to glass vials for injection into an ultra-performance liquid chromatography system coupled to a Q Exactive mass spectrometer in positive ionization mode (UPLC-MS), according to established protocols (Lapidot-Cohen et al., 2020). In any case, both for the primary and lipid metabolic datasets, standardized metabolite intensities (standardized by internal standard and weight) were then batch-corrected (ComBat), normalized by *glog* transformation and autoscaled (mean-centered and divided by the standard deviation of each variable) using MetaboAnalyst v5.0(Pang et al., 2021). The results for metabolites are reported following the standards suggested in Fernie et al. (2011).

### Fruit biometry

Twelve fruits were sampled from each parental and hybrid genotype. In addition, as interspecific hybrids showed different morphology and large increases in fruit size when they received pollen from parents, we decided to include them in these analyzes. The fruits were cut longitudinally and were digitized in a scanner (HP Scanjet G2410). In some cases, the fruits had the biometric contours adjusted manually in order to correct points of non-detection in the image. The biometric determinations were carried out using Tomato Analyzer v4.0, and 32 parameters were measured as previously described (Rodríguez et al., 2010).

### Estimation of heterosis

The experiments were carried out in a completely randomized design, always with the parents being cultivated together with their hybrids in their respective growing season. Each set of phenotypic traits were subjected to analysis of variance; in those cases where the null hypothesis of equal means was rejected, we used the Scott-Knott test with alpha=0.05 to identify the significant differences. Heterosis calculations were performed for the main parameters evaluated, using the following equations (Barth et al., 2003):

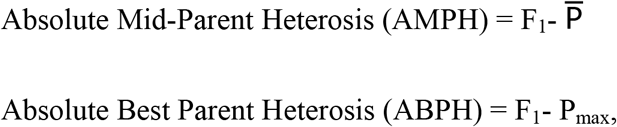

where: F_1_: average value of the phenotypic trait from the F_1_ hybrid; 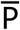: average value of the phenotypic trait of the two parental genotypes; and P_max_: Average value of the phenotypic trait from the best parent.

In addition, heterosis relative values were calculated as:

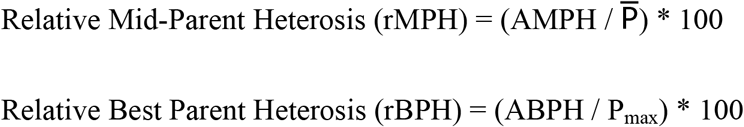

The reciprocal effect for the HAB × JAL, JAL × IKE and HAB × BIQ hybrids was determined as the difference in a given parameter between the reciprocal hybrids, e.g.: (HAB × JAL) - (JAL × HAB). The absolute values of heterosis and reciprocal effect were subjected to ANOVA and subsequently subjected to Scheffe’s test in order to verify the significance of each value individually.

## Acknowledgements

AZ acknowledges a CAPES/Alexander von Humboldt Foundation fellowship. We thank Prof. Fábio DaMatta (UFV) and Prof. Lázaro Peres (USP) for valuable input in the project.

